# Investigating action topography in visual cortex and deep artificial neural networks

**DOI:** 10.1101/2025.08.05.668643

**Authors:** Davide Cortinovis, Nhut Truong, Hans Op de Beeck, Stefania Bracci

## Abstract

High-level visual cortex contains category-selective areas embedded within larger-scale topographic maps like animacy and real-world size. Here, we propose action as a key organizing factor shaping visual cortex topography and assess the ability of topographic deep artificial neural networks (DANNs) in capturing this organization. Using fMRI, we examined responses to images of body-parts and objects with different degrees of action properties. In left lateral occipitotemporal cortex, we identified a topographically-organized action gradient, with overlapping activations for bodies, hands, tools, and manipulable objects along a dorsal-posterior to ventral-anterior axis, culminating at the intersection of body parts and objects exhibiting higher action properties. Multivariate analyses confirmed action as a crucial organizing principle, while shape and animacy dominated ventral occipitotemporal cortex and DANNs, which exhibited no action-based organization. Our proposed action dimension serves as a further organizing principle of object categories, advancing understanding of visual cortex organization and its divergence from DANN-based models.

## Introduction

Topography – the systematic, spatial organization in which neurons (or voxels) with similar functional properties are located near one another in the cortex^1^ – is ubiquitous throughout the cortex, from the retinotopy and pinwheels of primary visual cortex^2^ to the complex somatotopic organization of body parts in the so-called motor homunculus in M1^3^. In occipitotemporal cortex (OTC), a topographic organization of functionally selective areas has been shown, with areas responding preferentially to ethologically-relevant categories such as faces, body parts, words, and scenes^4,5^, mirrored along the ventral and lateral OTC^6^, and forming a consistent spatial arrangement across participants^7^.

Several accounts have tried to explain this organization by the role of different features that map object space onto the two-dimensional cortical sheet, leading to the emergence of functionally selective areas. These features span from low-level principles like eccentricity^8,9,10^, to mid-level properties (e.g., curvature^11^, aspect-ratio^12,13^, texture^14^), and to semantic principles like animacy^15^ and real-world size^16^. Some of these dimensions appear to be repeated across ventral and lateral OTC, explaining the mirrored organization of category-selective areas^17,18,19^. Remarkably, the representational space of higher-level layers in DANNs trained on object recognition captures the same object dimensions observed in the visual cortex (e.g., animacy^20^, aspect-ratio^12^ – but see^21^ – shape^22^, real-world size^23^). Moreover, topographic DANNs – architectures that incorporate biologically inspired spatial constraints^24,25,26^ – develop category-selective responses (e.g., for faces, bodies, and scenes) that mirror the topographic organization found in the visual cortex.

Notably, accumulating evidence suggests that despite the fact that lateral and ventral OTC show a similar mirrored object topography, their underlying representational space might be better explained by different object dimensions^27,28^. For instance, the left lateral OTC shows sensitivity to categories characterized by their action-related properties such as hands and tools^29,30,31^, whose underlying selectivity is spatially adjacent to, and partially overlaps with, one another^32^. Hands and tools differ in many visual and semantic properties, such as their shape and animacy; eccentricity and real-world size accounts also cannot explain this pattern of results as this effect does not extend to other object categories sharing similar eccentricity or real-world size^33,34^. Instead, this evidence suggests that another dimension plays a role in shaping the topographic organization of visual cortex object space: action^33^.

The present study aims to investigate the principles underlying the organization of functionally selective areas, with a focus on how behaviorally relevant action properties of objects shape the spatial organization and content of representations in ventral and lateral OTC. We conducted an fMRI experiment where participants viewed images^35^ of body parts and objects varying in their degree of action properties.

Using univariate and multivariate analyses on fMRI data, along with representational predictions based on human similarity judgments, we tested how action dimensions interact with other proposed dimensions and compared results in human visual cortex with DANNs. Our results show a dissociation between ventral and lateral OTC in both topography and representational space. Action—alongside shape and animacy—emerged as a key principle explaining the arrangement of categories in lateral OTC, while animacy best explained topography and representational content in ventral OTC and in DANNs, which in turn did not show any action-related organization. These results demonstrate that action is a fundamental organizing dimension of OTC, and that further developments are necessary for current computational models to fully capture both topography and function of high-level visual cortex.

## Results

To investigate how action-related properties influence object topography in visual cortex, we designed a stimulus set organized along two dimensions: animacy (body parts vs. inanimate objects) and action. Specifically, the three inanimate categories vary along two action-related properties: action effector and graspability (Figure 1). Tools are both action effectors and graspable; manipulable objects are graspable but not effectors; and non-manipulable objects are neither effectors nor graspable. The three body parts also differed in action relevance: low for faces, higher for bodies, and highest for hands. Action-related properties for all categories were behaviorally validated (see methods for details).

**Figure 1.**
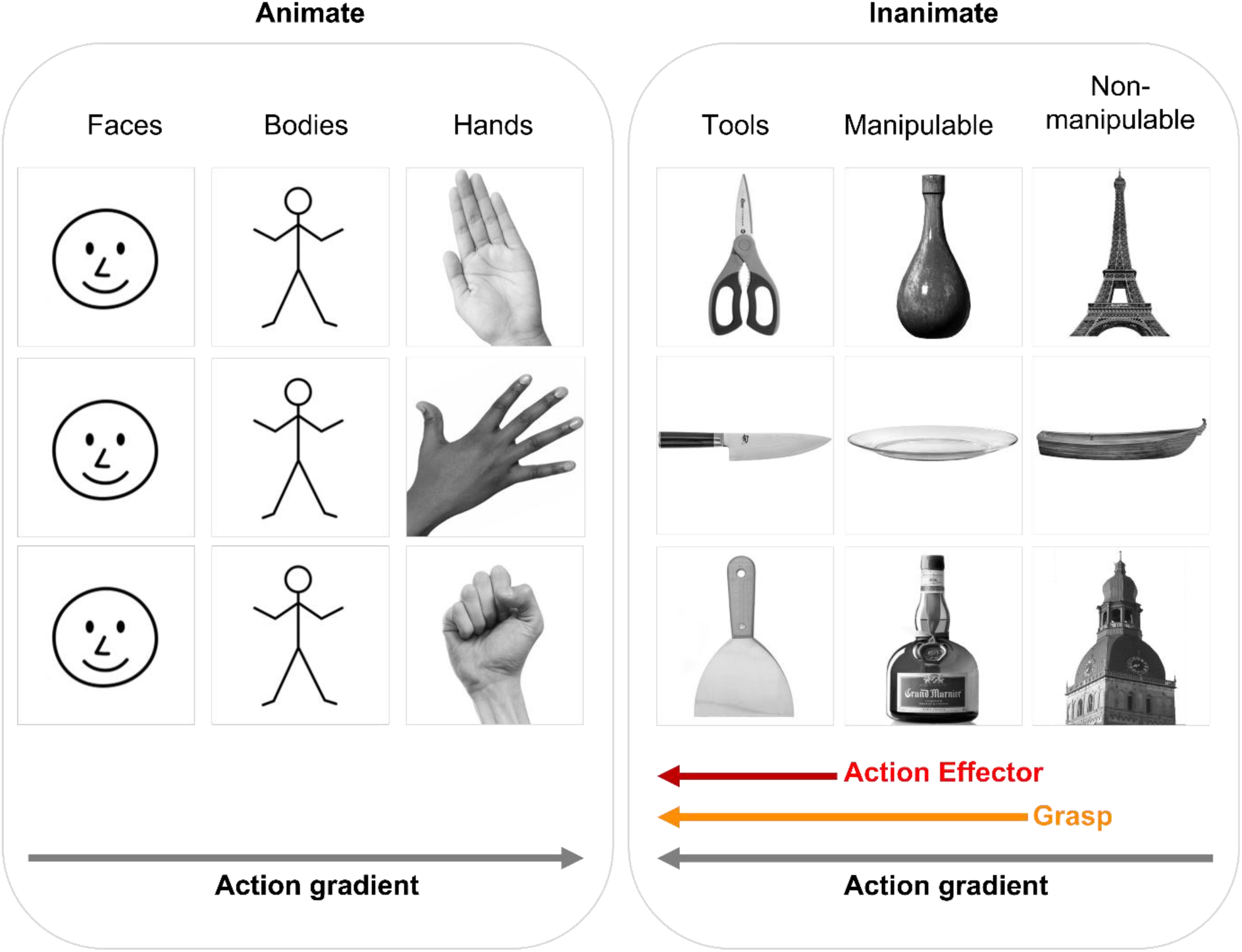
Stimulus set. Images were divided into 6 categories varying along two dimensions, animacy and action. For inanimate objects, action was characterized by two properties, end-effector and graspability. The three inanimate objects were matched for visual shape and orientation, to avoid confounds based on the overall shape (e.g., the elongation) of the stimuli. Stick figures were used in place of images of people.

To investigate the degree to which animacy and the two properties of the action dimension can predict the object topography in visual cortex, we combined univariate (e.g., functional profile, overlap analysis) and multivariate analyses of fMRI data to examine both the large-scale spatial distribution and the underlying representational content in lateral and ventral OTC. In parallel, we evaluated the ability of DANNs to capture this organization to assess where current models align with, or diverge from, biological systems.

### Action properties differentially shape object topography in ventral and lateral OTC

To investigate object space organization in ventral and lateral OTC (VOTC and LOTC, respectively), we first mapped the activation response for each category (versus all others, t > 3.5, p < .05 FDR corrected at the cluster level) onto the whole-brain surface (Figure 2a). Beyond replicating the known parallel organization of category selective responses in lateral and ventral OTC^36^, the whole-brain analysis confirmed a dissociation between the VOTC and LOTC in the left hemisphere (Figure 2a) based on the activation patterns for object classes with varying degrees of action-related information. Whereas in VOTC we found the typical medial-to-lateral animacy division with no overlap between animate and inanimate categories^7^, in LOTC we observed overlapping responses between animate and inanimate conditions with a different degree of action properties. From dorsal-posterior to ventral-anterior, we observed selective and partly overlapping activations for bodies, hands, tools, and manipulable objects, with a convergence and high degree of overlap for the animate and the inanimate categories characterized by the highest degree of action properties: hands and tools. The action-based organization was particularly evident when comparing activations of inanimate objects. Specifically, we found a consistent action-related gradient in LOTC, with a smooth transition across the cortical surface where the activation to object categories characterized by different action properties changes systematically according to the two action-related properties. This gradient was characterized by a large activation cluster for tools which are both action-effector and graspable, a smaller cluster for manipulable objects which are only graspable, and no significant activation for non-manipulable objects which are neither action effector nor graspable; the opposite pattern was observed in VOTC, with a larger cluster for non-manipulable relative to manipulable objects, which in turn revealed a larger activation relative to tools. The action-related topographic organization in LOTC was also observed at the level of individual participants, without spatial normalization or smoothing (see Figure 2c for an example participant). Unlike the left hemisphere, the right hemisphere did not show any action-related organization, as neither tool nor object selectivity were observed (see Supplementary Materials for right hemisphere results). In the remainder of the paper, all analyses refer to the left hemisphere.

**Figure 2.**
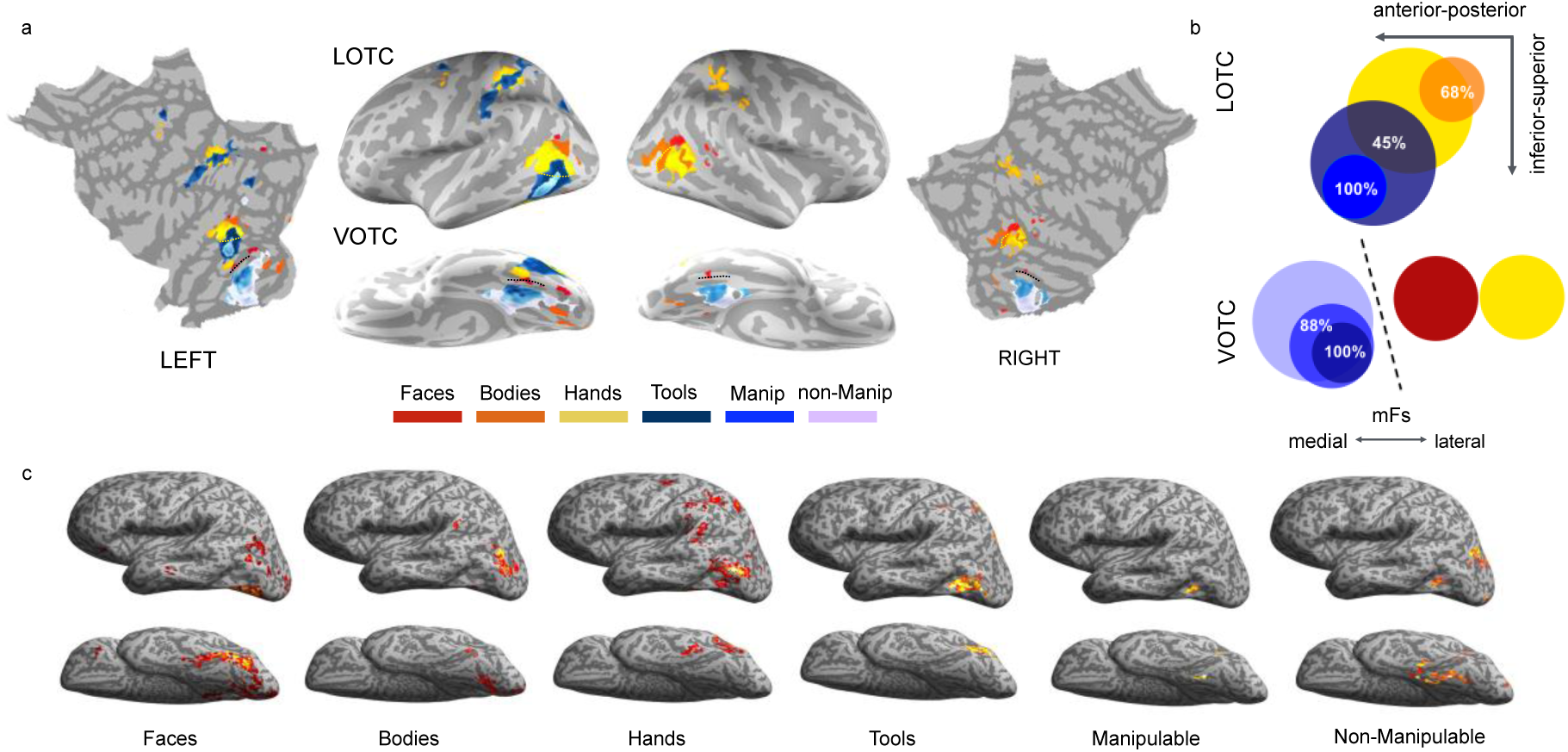
Action-related topography of occipitotemporal cortex. a) Whole-brain results. Response for each category (vs. all) was visualized on a freesurfer average brain surface using BrainSurfer (https://www.mathworks.com/matlabcentral/fileexchange/91485-brainsurfer), with a threshold of t > 3.5 (p < .05 FDR corrected at the cluster level), excluding activations within early visual cortex (approximately V1-V2-V3) to focus on the regions of interest in LOTC and VOTC. Color-coded dashed lines indicate overlap between activations. The black dashed line indicates the mid-fusiform sulcus. b) Category overlap visualization. The size of each circle represents the approximate size of the category-selective cluster in VOTC and LOTC in the left hemisphere. c) Single subject results on the unsmoothed native surface of one representative participant (t > 3.5, FDR cluster corrected at p < .05).

These results were further confirmed by the overlap analysis, which allowed us to further assess the spatial relationship between categories, with the underlying rationale that spatial proximity and overlap in the cortex suggest shared features^37^. We quantified the extent of activation overlap between each category by calculating an overlap index for each pairwise combination of regions, separately for the ventral and lateral OTC (see methods, Figure 2b). The index represents the number of voxels common between the areas, varying from 0 (no voxels in common) to 1 (the smaller area falls completely within the larger). In LOTC, from dorsal-posterior to ventral-anterior a large overlap could be observed between hands and bodies (0.68), between hands and tools (0.45), and between tools and manipulable objects (1.0, where manipulable objects fall completely within the larger tool cluster), but no overlap could be observed for the other combinations. On the contrary, in VOTC, no overlap could be observed between animate and inanimate categories, nor between faces and hands; inanimate objects, instead, presented a strong overlap with each other, with tools falling completely within the manipulable object cluster (1.0), and manipulable showing an extended overlap with non-manipulable objects (0.88), thus further confirming the opposite gradient in LOTC and VOTC for objects characterised by a different degree of action properties. A schematic visualization of category overlap is shown in Figure 2b.

To further characterize the spatial and functional profile of the different object topography observed in LOTC and VOTC, we plotted the beta values for each condition extracted from a series of partially overlapping spheres covering a broad region of visual cortex including a wide portion of ventral and lateral OTC from the parahippocampal cortex (PHC) to the transverse occipital sulcus (TOS) (see methods, and Figure 3a). The vector of ROIs analysis confirmed that from lateral to ventral OTC, the response profile for all inanimate objects follow a similar activation trend but with an opposite response strength based on action-related properties of objects: tools, which are both action effectors and graspable, show the highest response peak in LOTC and the lowest in VOTC; manipulable objects, which are graspable but do not serve as effectors, show the intermediate response in both LOTC and VOTC; and non-manipulable objects which are neither action effectors nor graspable show the lowest response in LOTC but the highest in VOTC (Figure 3a).

**Figure 3.**
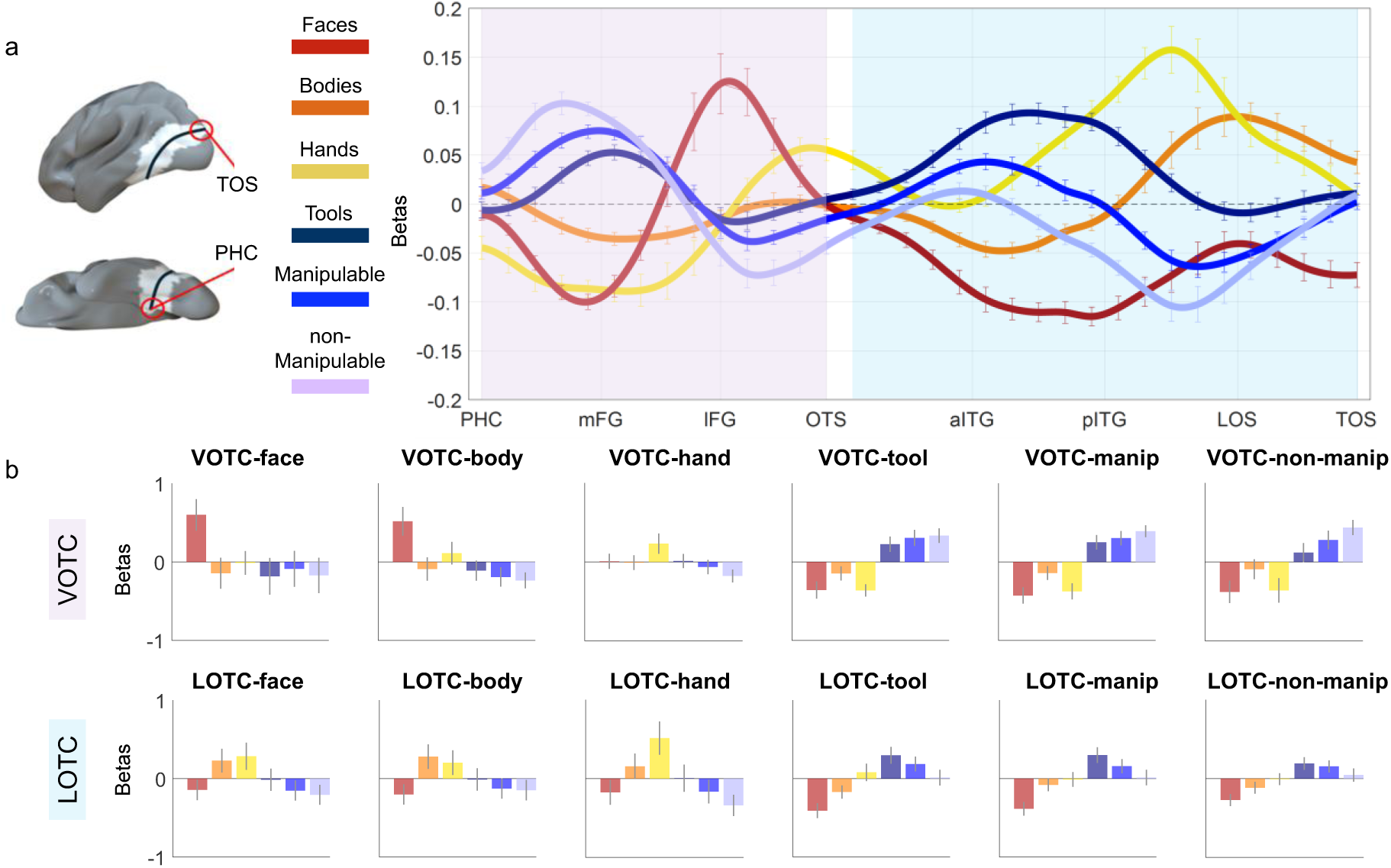
Distinct object topographies in lateral and ventral OTC. a) Vector-of-ROIs analysis. The vector was generated by fitting a spline (drawn black line) connecting the PHC and the TOS and passing through a set of anchor points which coordinates were based on classically defined category-selective areas (i.e., face, body, hand, object) from previous studies. Partially overlapping spheres (n = 34) were generated along this spline, and they correspond to the ROIs analysed. Standard univariate analyses were performed on each of the ROI (see methods for details), which are visualized in white with a surface projection using Surf Ice (https://www.nitrc.org/projects/surfice/). Normalized activation (against the average of all categories) is plotted for each category as a function of the position of the vector along the cortex. The x-axis corresponds to each sphere along the vector, with labels for major anatomical landmarks; the y-axis corresponds to the normalized beta values. The vector was broadly divided into a ventral component (pink shade) and a lateral component (light blue shade). b) Beta values are plotted for each category’s peak activation (one sphere) separately for the ventral and lateral left OTC. Error bars correspond to ± 1 SEM across subjects. PHC = Parahippocampal Cortex. mFG = medial Fusiform Gyrus. lFG = lateral Fusiform Gyrus. OTS = Occipitotemporal Sulcus. aITG = anterior Inferior Temporal Gyrus. pITG = posterior Inferior Temporal Gyrus. LOS = Lateral Occipital Sulcus. TOS = Transverse Occipital Sulcus.

Overall, these results indicate that the topography of objects in lateral and ventral OTC is driven by their different degree of action properties, as measured by their action-effector and grasp properties. To verify this, we plot the peak response (1 sphere) for each condition in ventral and lateral OTC (Figure 3b). These results confirm that, within the inanimate object cluster, tools elicit the highest activation across all three LOTC object peaks (*p* < .01 for all contrasts; corrected for n = 5 comparisons; *p <* .01). In contrast, non-manipulable objects elicit the highest response across all three VOTC object peaks (*p* < .001 for all contrasts), except in VOTC-tool where non-manipulable and manipulable did not differ from each other (*p* = .41). Hands elicit the highest activation in the LOTC animate peaks (LOTC-hand, LOTC-face) compared to all other object categories (*p* < .001 for all contrasts) except for LOTC-body where bodies elicited the highest response (*p* < .003 for all contrasts). Finally, whereas faces show the typical selectivity in VOTC (VOTC-face and VOTC-body: *p* < .001 for all contrasts), we also observed a small but selective cluster for hands in the occipitotemporal sulcus, located lateral to the face cluster, which shows significant higher activation for hands than for all other categories including faces and bodies (VOTC-hand: *p* < .001 for all contrasts). This region likely corresponds to the left counterpart of the fusiform body area^38^, a region that has been also called OTS-limbs^31^. Here, we report its selective activation for hands specifically and not bodies in general, thus confirming the possibility of dissociating the activation to hand stimuli from the one to whole bodies not only in lateral^29^ but also in ventral OTC (see also^36^).

Overall, these results support the conclusion that the “parallel” object representations in LOTC and VOTC encode distinct object properties, and specifically point to the presence of an opposite organization within ventral and lateral OTC, with the latter being sensitive to object categories that contains a different degree of action information, as indexed by the consistent topographic organization for objects and body parts with different action-related properties and convergence between inanimate (tools) and animate (hands) categories that share effector properties.

### Topographic DANNs successfully mimic animacy division in VOTC but fail to replicate action-based topography in LOTC

The above results show that the lateral and ventral OTC are characterized by a different topographic organization: whereas in VOTC the animacy of objects strongly drives the organization of representations giving rise to the well-documented animacy division, in LOTC the topographic organization is driven by the degree of object action properties with a gradient from posterior-superior to anterior-inferior. Here, we test whether topographic deep artificial neural networks (TDANNs), a type of computational model developed to capture the topographic organization of ventral visual cortex^26^, can mimic the action-related organization observed in lateral OTC. TDANNs allow testing whether a model designed to capture general topographic organization as a by-product of minimizing wiring-length^39^ can account for object topography in visual cortex, thus suggesting that brain-like representations and their spatial organization can co-emerge with biologically inspired spatial constraints.

The network architecture was based on a ResNet-18 backbone, pre-trained with a self-supervised contrastive-learning object recognition task^40^. We tested five different random initializations of the network’s weights. We fed the networks with the images from our experiment and extracted the activation maps for each topographic layer, selecting the last “VTC-like” layer for further analyses (consistent with^26^). A unit was defined as selective if its response for a specific category passed a set threshold (defined as *t* > 3.5, with a contrast of category > all). This uncorrected threshold was chosen for visualization purposes only (Figure 4a). The subsequent functional selectivity analysis was performed on the first 50 most selective units. To investigate whether TDANNs replicate the topography and functional profile of category activations in visual cortex, we visualized their respective spatial distribution in the simulated cortical space and plotted the activation profiles for the 50 most selective units per category. Results are shown in Figure 4. Despite some variations between the five initializations – especially in the clustering’s strength – two main findings could be observed (Figure 4a): first, in all networks, units selective for animate and inanimate objects formed separate clusters, such that when a unit responded to a body-part it did not respond to an inanimate object and vice versa; second, no organization based on action properties was observed. Specifically, tools and hands did not activate the same units, and no smooth overlap based on action properties was found among the three object categories.

**Figure 4.**
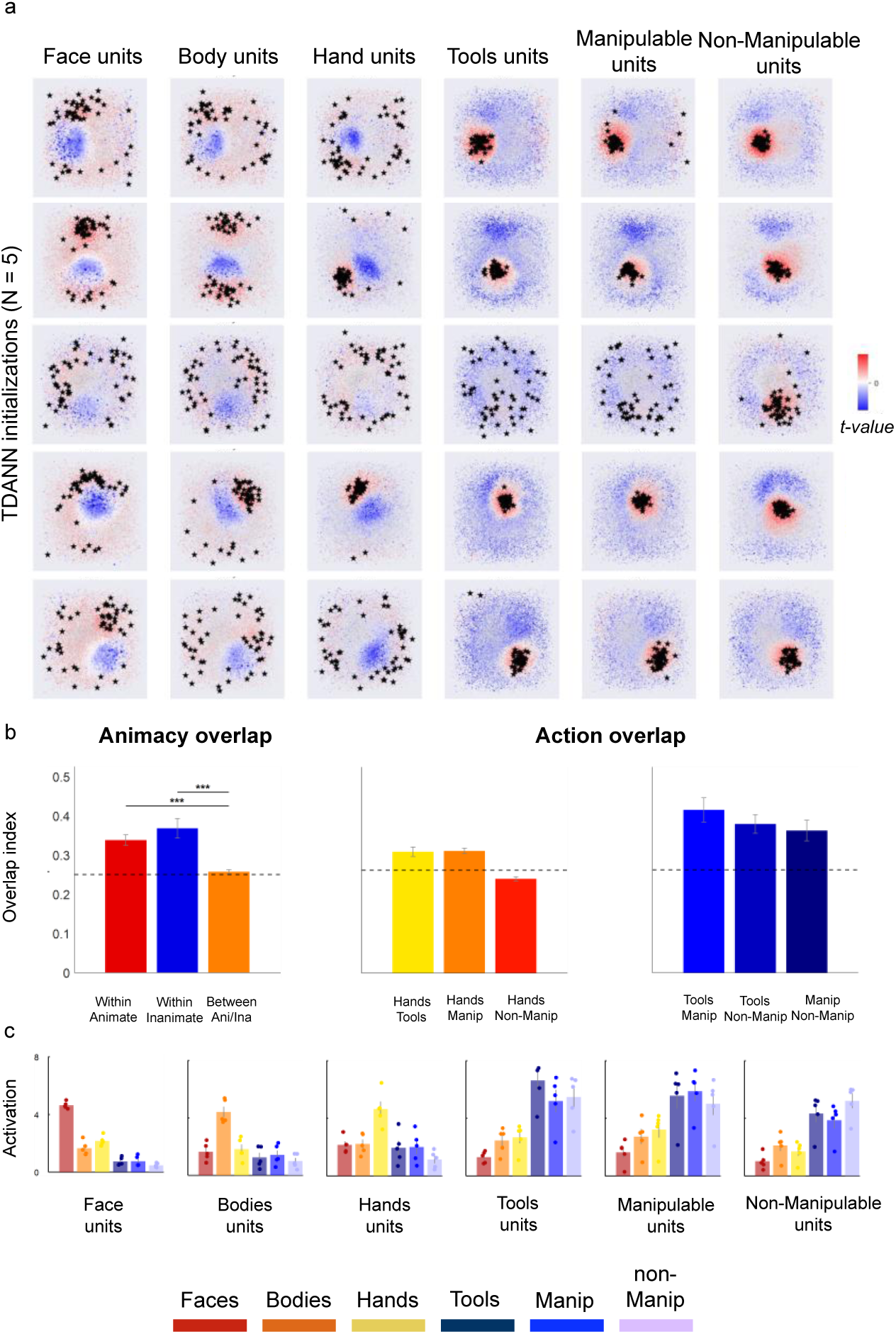
TDANNs replicates animacy but not action-related organization of OTC. a) Spatial distribution of each category (as defined by t-values) on the simulated cortical space of the VTC-like layer of five random initializations of the TDANN. Rows correspond to each of the five initializations. Stars represent the location of the top-50 most selective units for that category. Category-selective units (positive t values) are shown in red while units not selective for that category (negative t values) are shown in blue. b) Overlap analysis. Stars represent statistical significance (*p* < .001) computed across the five random initializations of the TDANNs, tested with 10000 permutations. Error bars correspond to ± 1 SEM across the random initializations. Black dashed line represents baseline (overlap of 0.5 means no correlation between the presence of two categories). c) Selectivity profile of the top-50 most selective units for each category, based on the activation of the VTC-like layer (as in a). Each data point represents the value from a single TDANN initialization.

To quantify these observations and compare TDANNs with brain results, we performed the overlap analysis (as in^26^). Specifically, we measured the co-occurrence of units selective for each category by using an overlap score ranging from 0 (the presence of one category always predicts the absence of the other) to 0.5 (no relationship) to 1 (perfect co-occurrence). Statistical significance was tested via 10000 permutation tests. Results (Figure 4b) confirmed significant overlap within animate (score: 0.68, *p* < .001) and inanimate (score: 0.74, *p* < .001) categories relative to the between-category overlap (animate-inanimate, score: 0.51). In other words, units that responded to a body part or an inanimate object also responded significantly to other categories within the same superordinate class. Second, the overlap score between action effector categories such as hands and tools (score: 0.59) was not significantly higher than the overlap between hands and other manipulable objects (score: 0.594, *p* = .37), as well as the overlap between tools and manipulable objects (score: 0.79) was not significantly larger relative to the overlap between tools and non-manipulable objects (score: 0.72, *p* = .24), nor relative to the overlap between manipulable and non-manipulable (score: 0.72, *p* = 0.33) thus, showing no action-related organization in TDANNs.

Visual exploration of Figure 4a suggests that, in addition to the separation between animate and inanimate categories, there seem to be additional differences in the organization of categories. Specifically, whereas the spatial distribution of units selective for the different body parts seem a bit scattered around, the inanimate objects mostly activated a similar portion of the cortical space. To investigate the functional profile of the TDANN units, we extracted the activation profiles for the 50 most selective units for each category and plotted the results (Figure 4c shows results averaged across the five initializations). Here, the focus was not on unit selectivity per se (e.g., do tool units respond to tools more than all other categories) but rather the degree to which a unit that responds to one category also responds to other categories (e.g., do tool units respond to other categories as well?). Overall, the results show that while a certain degree of category-selectivity could be found for the different body-parts, as different units selectively activated for each body part independently from the other body parts, the top-units for each inanimate object category responded to the other inanimate objects to a similar degree. Indeed, the selectivity of units chosen based on their response for faces, bodies, and hands was significantly higher for their preferred category compared to all other categories (for all contrasts, *p* < .001; permutation test n = 10000). In contrast, units selected for their response to tools, manipulable and non-manipulable objects did not differ in selectivity across other inanimate object categories (for all contrasts, p > .05), while being more selective for their preferred category than for the animate categories (for all contrasts, *p* < .001; permutation test n = 10000). Thus, similar to what we observed in visual cortex, TDANNs units that respond to one inanimate object category do also respond to the other inanimate object categories, but differently from human VTC, we did not observe any differential response gradient from high to low (tools > manipulable > non-manipulable) as observed in LOTC or from low to high (non-manipulable > manipulable > tools) as observed in VOTC. Finally, differently from visual cortex, units that respond to tools did not seem to activate hand units, thus confirming results from the TDANNs overlap analysis.

Overall, these results show that TDANNs primarily distinguish between animate from inanimate objects, with additional functional selectivity for individual body-parts, and a weaker, or absent, distinction among inanimate object categories. These results mirror the pattern of overlap found in VOTC, which also showed a separation between animate and inanimate object categories, with further clustering for hands and faces. However, no action gradient organization, as found in LOTC, could be observed in TDANNs.

Altogether, these analyses on networks implementing biologically-inspired topographic constraints reveal their ability to capture visual features important to distinguish animacy and to capture– to a certain extent – the selectivity for body-parts, but cannot capture the action-related organization observed in visual cortex.

### VOTC and LOTC support distinct object feature spaces

Our results reveal a different object organization in LOTC and VOTC and that TDANNs are able to capture only part of visual cortex topographic organization. Next, we employ multivariate analyses to further investigate what properties underlie this object space. Specifically, we use representational similarity analysis (RSA^41^) to investigate how the action and animacy dimensions relate in both visual cortex and DANNs. We created three models, each reflecting a distinct dimension: the action model capturing action-related information for each object category; the animacy model capturing the body-parts\inanimate objects divide; and the shape model capturing the average aspect-ratio of each category (see methods), added to account for visual properties relevant in OTC^12,13^. The animacy and action models were generated from participants who judged a random subset of stimuli (n = 36) on each dimension (see methods). The models were orthogonal: animacy vs. action-effector (r = – 0.08); animacy vs. shape (r = 0.08); action-effector vs. shape (r = –0.16). Dissimilarity matrices (Figure 5a) support our predictions: the animacy model clearly separated body-parts from inanimate objects; the action-effector model showed a graded continuum: as the action-related properties of body parts and objects increased, their correlation strengthened.

**Figure 5.**
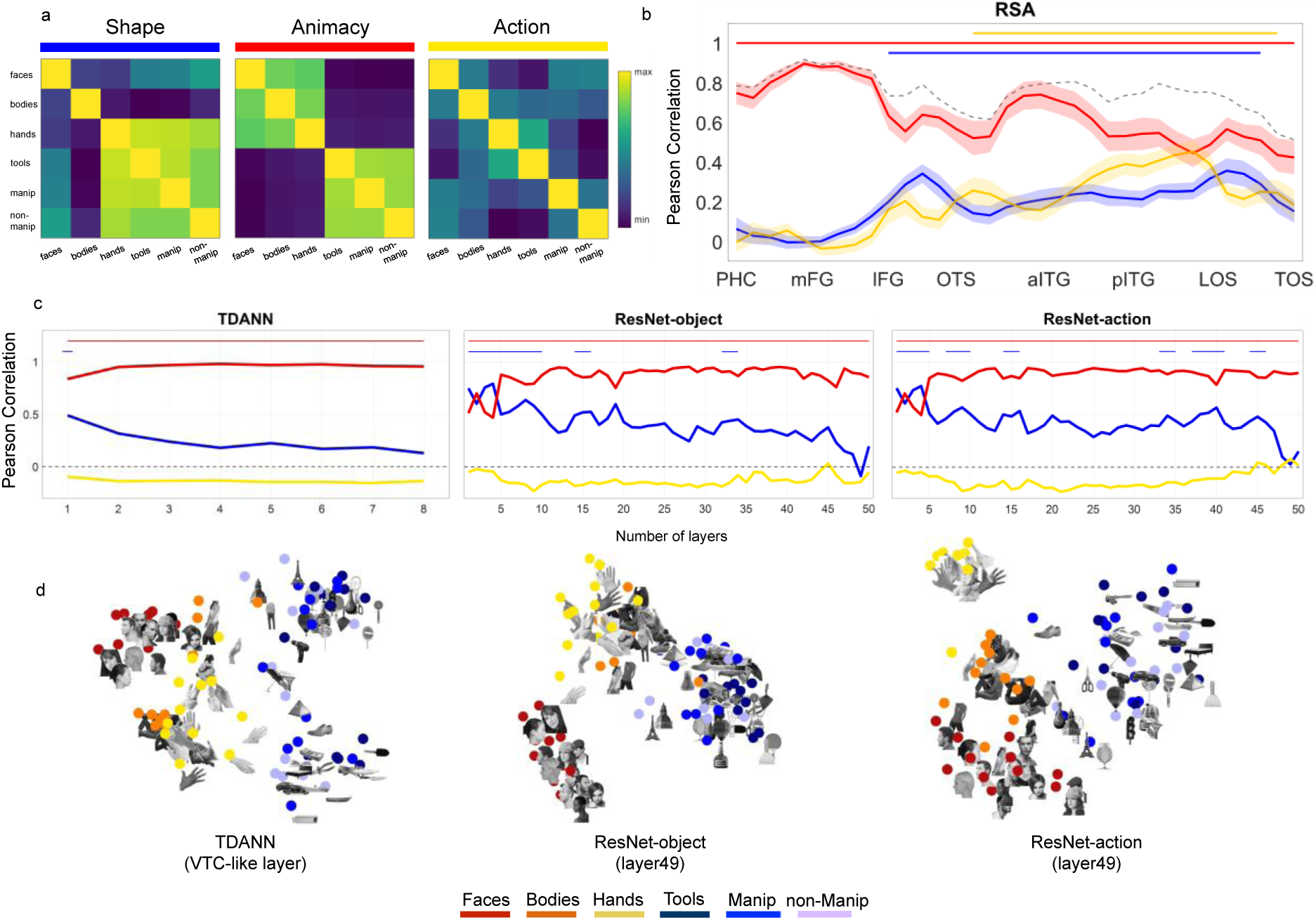
The distinct role of action and animacy in visual cortex and DANNs. a) RSA Models: the shape model captures the aspect-ratio of the stimuli, whereas the animacy and action models are based on behavioral ratings (see methods). b) Vector-of-ROIs RSA results. The dashed line represents the noise ceiling boundary which indicates the highest best possible fit to the neural data that a model can achieve given the noise in the data. Horizontal lines indicate statistical significance (vs. baseline) for each model (p < .0014 Bonferroni corrected for n = 34 comparisons). c) RSA results for the three DANNs. Color-coded lines on top of each graph indicate the layers where each model reach statistical significance relative to baseline (p < .001; 10000 permutations). d) MDS for the DANNs (ResNet-Object and ResNet-Action) last convolutional layer (layer 49) and the TDANN VTC-like layer. Results for the TDANN refers to one of its initializations.

We assessed how these dimensions are represented across lateral and ventral OTC, by correlating neural activity patterns in each vector-of-ROIs sphere with the three models (Figure 5a). Results showed that while animacy was strongly represented across the entire swath of cortex and reached the noise ceiling in ventro-medial regions of OTC, the action dimension reached its highest peak within LOTC, specifically between posterior ITG and LOS and its lowest peak in VOTC, in correspondence of the highest peak for animacy. Interestingly, throughout both ventral and lateral OTC, the effect for object shape closely followed the trend of the action model, suggesting that regions encoding action-related properties of objects also represent their shape properties. To quantify this trend, we perform pairwise correlations between the effects of each model along the vector. Results confirmed that shape and action did indeed show a small but significant correlation along the vector (*r* = 0.18, *t*_(17)_ = 3.2, *p* = .0044; for all RSA results, correction for n = 3 comparisons; *p* < .016). On the contrary, a significant negative correlation was observed between the action and the animacy models (*r* = -0.4, *t*_(17)_ = -8.2, *p* < .001), whereas no correlation was found between shape and animacy (*r* = -0.05, *t*_(17)_ = -1.2, *p* = .24).

We performed the same RSA analyses in the TDANNs and in two non-topographic models, both based on the ResNet-50 architecture but trained with different task objectives: object recognition with ImageNet^42^ (ResNet-object) and action recognition with Moments-in-Time^43^ (ResNet-action). This allowed us to test whether training objectives influence the networks’ representational space and whether action recognition training improves the representational correspondence with LOTC.

The RSA analysis performed in DANNs revealed different results. Across all networks, regardless of architecture (topographic or non-topographic) or training task (object or action recognition), animacy was the dominant dimension, highly significant throughout the network hierarchies and outperforming other models in most layers (Figure 5c). Shape was the second-best model, with high correlations along the networks’ hierarchy dropping in the final layers, in line with previous reports^22^. The action model never reached significance in any layer or model. Furthermore, differently from what we observed in visual cortex, action and shape were not significantly correlated across DANNs’ layers (Pearson *r* = –0.14; p = 0.34). Together, these results show that neither training task is sufficient to produce a brain-like action-related organization in the networks.

To further inspect the DANNs feature space, for each model we projected the dissimilarity matrix of the last convolutional layer (layer 49) of the two ResNet and the VTC-like layer of the TDANN into a two-dimensional plot by using multidimensional scaling (MDS; Figure 5d). Confirming the RSA results, the animacy division appears to be the main dimension emerging in the representational space of all DANNs with no evidence for any action gradient. In addition, an effect of shape was observed in the arrangement of inanimate objects. That is, differently from body parts, which show some clustering based on category, objects that by design were matched for shape, show an arrangement based on visual properties such as aspect-ratio and orientation.

### Lateral OTC represents action-effector and (to a lesser extent) grasping properties of objects

Up to now, we have shown that distinct object dimensions are represented in ventral and lateral OTC. Here, we further characterise the specific action-related properties underlying this object space. To this aim, we calculated two indices derived from the correlational matrices obtained with multivariate analysis (see methods): the action-effector index and the grasp index. The indices measure distinct properties of the object categories, specifically the possibility of an object to be an end-effector (the action-effector index), which differentiates tools (e.g., a pair of scissors or a knife) from other graspable objects (e.g., a bottle or a glass) and is shared between hands and tools, and the possibility of an object to be grasped (the grasp index), which differentiates manipulable objects from large non-manipulable objects that cannot be grasped (e.g., a building or a vehicle). The action-effector index was calculated by taking the correlation between each body-part with tools and from that subtracting the correlation between each body-part and manipulable objects; the grasp index was calculated by taking the correlation between each body-part with manipulable objects and from that subtracting the correlation between each body-part and non-manipulable objects (see methods). Results are shown in Figure 6.

**Figure 6.**
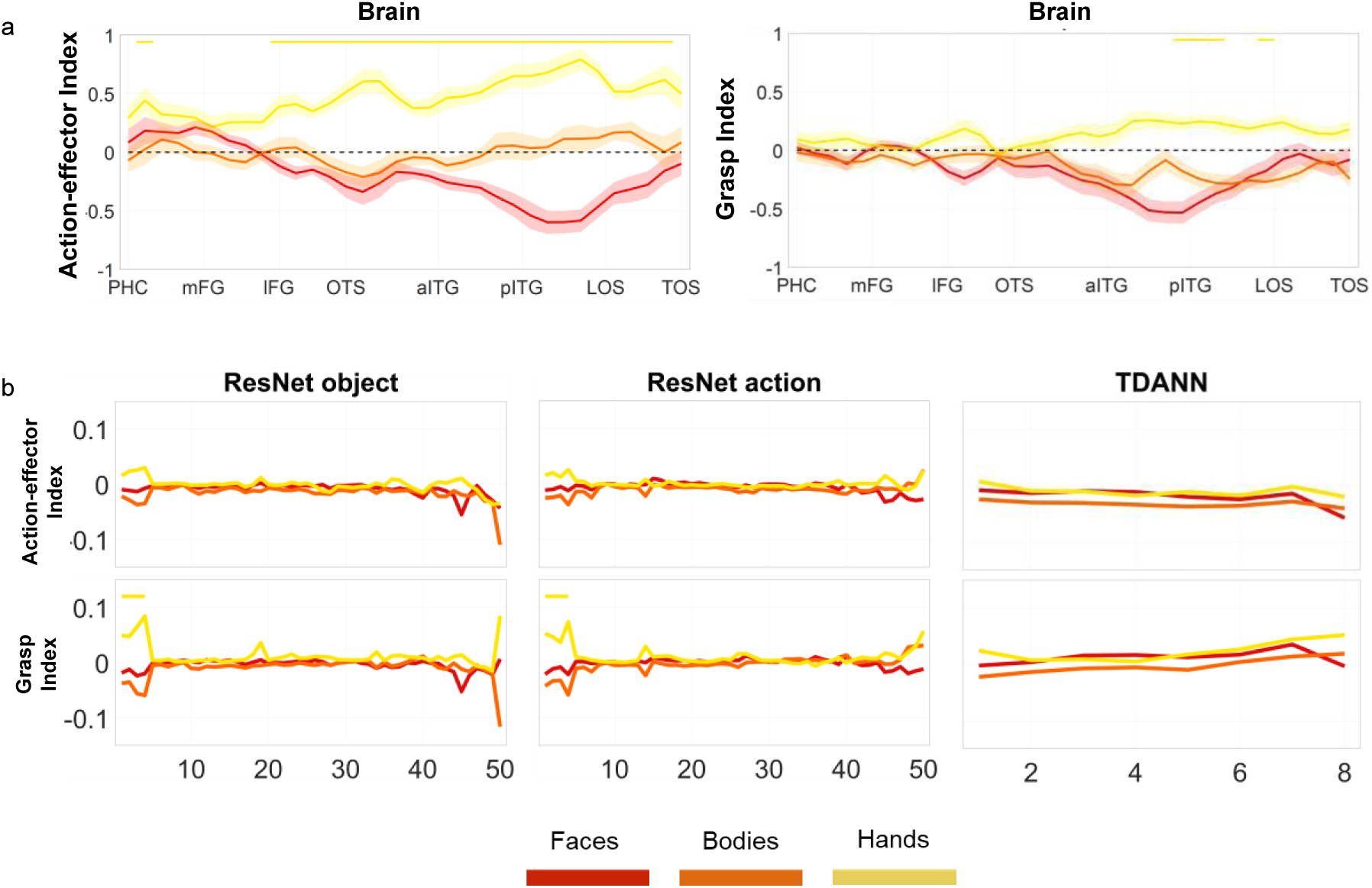
Index analysis. a) Vector-of-ROIs action-effector index (*right)* and grasp index (*left*). b) Action-effector index (top) and grasp index (*bottom*) for the three artificial networks tested. Color-coded lines at the top of each plot indicate spheres along the vector or layers where each index reached significance, corrected for the number of spheres (n = 34; *p* = .0014) and for n = 10000 permutations (*p* < 0.001) for brain and DANNs, respectively.

This analysis revealed that the driving factor underlying the object space in LOTC is the action-effector property of objects, followed by a smaller but significant effect for object grasp. More specifically, the action-effector index shows that across the whole LOTC, hands are strongly associated with objects that are characterized by effector properties, such as tools, compared to other manipulable objects which share graspable properties with tools but do not serve as action effectors (Figure 6a, left). This effect is specific for hands, as whole bodies do not show the same pattern and faces even show a negative index (which indicates higher correlation with objects that are not action-effectors). These results show that while the action-effector effect is present throughout most LOTC, its strength follows closely the response profile of hands, suggesting that univariate hand-selectivity supports an object space with one of the main dimensions being action-related. To directly test this relationship, we computed the correlation between the effector index and the activation for the different object categories along the vector-of-ROIs. Throughout the vector, the effector index was significantly correlated with the hand’s response profile (*r*_(17)_ = 0.38; *t*_(17)_ = 4.46, p < .001; correction for n = 6 comparisons; *p* < .0083) but not with the response profile for either faces, bodies, or tools (faces: *r*_(17)_ = -0.09; bodies: *r*_(17)_ = 0.08; tools: *r*_(17)_ = 0.032;) and it was negatively correlated with the response profile for manipulable and non-manipulable objects (manipulable: *r*_(17)_ = -0.2; *t*_(17)_ = -3.61, p = .0022; non-manipulable: *r*_(17)_ = -0.28; *t*_(17)_ = -4.7, p < .001).

The grasp index (Figure 6a, right) reveals a smaller but significant effect in some regions of LOTC, showing that hands are also associated with manipulable objects more than to non-manipulable objects. This effect was not observed for bodies and faces. Confirming the other analyses, no significant grasp index was found in VOTC. Finally, in line with the weaker grasp-related effect, only a modest relationship was found between univariate selectivity for hands and the grasp index (hands: r = 0.22; p = .031) which however did not survive correction for multiple comparisons (n = 6; p > 0.0083).

Although DANNs do not show any action properties, for completeness and to test possible similarities or differences with visual cortex we calculated the action-effector and grasp indices for all layers and DANNs (Figure 6b). In agreement with the above results, no network shows either action-effector or grasp effects; the two indices did not reach significance (*p* > .05) at any stage of the hierarchy of any of the networks, except for a small effect in the first four layers of both non-topographic networks for the grasp index.

## Discussion

Our study identifies action as a fundamental dimension shaping the topographic organization of the visual cortex. We demonstrate that the left lateral occipitotemporal cortex (LOTC) exhibits a dorsal-posterior to ventral-anterior gradient where body parts and inanimate objects are topographically organized based on their action-related properties. The combination of action effector and graspability contributes to explain the spatial organization of voxels that show a preferential response to bodies^42^, hands^29,36,45^, tools^46^, and manipulable objects^47^. While DANNs replicate aspects of ventral stream organization (e.g., animacy), they entirely lack the action-related topography observed in lateral OTC. Together, our results show that the action dimension is an important organizing principle of lateral OTC and highlight remaining gaps between biological and artificial systems.

Previous work emphasised how the combination of multiple object dimensions and principles may result in the topography-by-selectivity that is observed in high-level visual cortex^7,48,49,50,51,52,53,54,55,56,57^, with proposals stressing the role of shape, animacy, and real-world size^16,18,58^, among others. Previous studies have already shown the relevance of action in explaining aspects of LOTC object space^27,59,60,61^. For example, overlapping responses in left LOTC between tools and hands, or tools and graspable food might reflect shared end-effector properties^32^ and action-related affordances^37^. Our results are in line with these previous findings and lift them up to a whole new level by revealing that a large-scale topographic organization is responsible for these earlier findings. More specifically, this approach enables us to move beyond post-hoc interpretations of visual cortex category organization (e.g., faces in lateral FG, tools in medial FG), allowing us to generate novel predictions about the spatial organization of new object categories – to be tested in future experiments – that share similar action-related features. Based on where these categories fall within a multidimensional feature space, we can predict their alignment within the topographic layout of OTC. For instance, as food items share grasping properties with manipulable objects, and are not action effectors, we expect them to map along the same action-based dimension and partially overlapping with manipulable objects, but not with hands.

Furthermore, we demonstrate that lateral and ventral OTC represent different object features, with their topographic organization exhibiting opposing response patterns that depend on the degree of action properties associated with objects. In left LOTC, the action-based topography culminated at the intersection between animate (hands) and inanimate (tools) as both being end-effectors. Dorsally and posteriorly, hands overlap with bodies and inferiorly and anteriorly, tools overlap with manipulable objects which share with tools grasping properties but not end-effector properties. This organization is consistent across participants (even in unsmoothed, native surface) and cannot be explained by differences in object size or shape as tools and manipulable objects are matched for real-word size and all object categories are controlled for their overall shape. The opposite object pattern can be observed in VOTC, with higher and more extended activation for non-manipulable than manipulable objects, and tools being embedded within the manipulable object cluster in medial VOTC. These findings challenge views that tool representations in VOTC reflect action-related properties^30^, suggesting instead that they encode general object features – such as surface properties^62^ or weight^63^ – shared across manipulable and non-manipulable objects to support recognition of inanimate objects in general rather than tools specifically^64,65^.

The opposite activation patterns observed in ventral and lateral OTC aligns with the proposal of a third lateral pathway dedicated to (inter)action recognition^27,28,66,67^ (see^68^ for a critical discussion). The studies characterizing this pathway have proposed a posterior-to-anterior organization, from perceptual to conceptual action-related and a medial-to-dorsal organization, from inanimate to animate processing and from transitive to social actions^27,28,69,70^. In this framework, the anatomical location of the LOTC action-based topography fall within a posterior and inferior region of the lateral visual pathway, suggesting their contribution to perceptually based action-related representations of objects.

But what is the origin of this action-based dimension? Although our experiment does not directly address this question, two alternatives might be considered. First, the action dimension might be perceptual in nature: for instance, hands and commonly used tools often visually appear together, which may explain why they are closely mapped in LOTC. According to the principle of minimizing wiring cost, which shapes known organizational patterns in both visual^1,39^ and motor cortices^71^, such visual co-activation may promote the proximity of hand and tool populations in LOTC. Alternatively, this dimension might be tied to motor experience with tools (e.g., learned associations between hands and tools during object interaction) reflecting how we engage with objects through action (but see^34^). Supporting this view, evidence shows that LOTC is active not only when viewing body parts or tools, but also during actual movements^45,72^. It is also plausible that multiple constraints might play a joint role in the emergence of this action-based topography, originating both from bottom-up visual factors (e.g., visual statistics) and top-down factors (e.g., behavioural goals) to ultimately represent object properties useful to support behaviour^49,53^.

Interestingly, studies have found that areas within the lateral visual pathway shows higher sensitivity to dynamic than static stimuli^73,74^. While the choice of static stimuli in the current study allowed us to have higher control on possible confounding variables (i.e., shape), future studies may employ dynamic stimuli such as short video clips of people performing actions that may not only replicate but even extend the relevance of behaviourally-relevant properties in explaining the object space in LOTC^75^.

Univariate and multivariate results revealed interesting couplings between object dimensions in visual cortex. Notably, object action and object shape representations were closely intertwined in lateral OTC, offering key insights into the functional organization of high-level visual cortex. The coupling of shape and action in lateral OTC highlights how object shape directly informs interaction potential. For instance, elongation – a mid-level shape property which characterizes most tools – is known to drive responses in tool-selective cortex^76^. Critically, however, our results go beyond these intrinsic associations between object category and shape^12,13^: even after controlling for shape, we observed robust action-shape coupling in lateral OTC, demonstrating that shape and action are distinct yet interacting dimensions.

DANNs results revealed both convergence and divergence with the functional and spatial organization of the visual cortex. Prior studies using topographic artificial neural networks^24,25,26^ or self-organizing maps^77,78,79^ have shown that principles like minimization of wiring length yield emergent macro- and mesoscale structures resembling those in visual cortex, including clusters for faces, bodies, scenes, and objects, and large-scale gradients of animacy and real-world size. Here, we confirm that while these networks capture the large-scale clusters based on animacy, and to a certain extent the category clusters for faces, bodies and hands, they could not capture the action-based object topography and the category clusters for the three inanimate object categories.

This failure may stem from DANNs’ reliance on mid-level visual features—such as shape and texture—that often correlate with object category in natural datasets. While this works well for animate categories (possibly because of curvature features^80^), it breaks down for inanimate categories when visual features are controlled, as in our study. In these cases, DANNs default to encoding lower-level properties like orientation or aspect ratio, leading to weak category-specific clustering for inanimate objects (Figure 5c-d). Thus, a tight control of visual features is especially important when comparing visual cortex and DANNs, as the two systems may represent objects in an apparent similar way but actually use different visual features that are confounded in the natural environment or uncontrolled stimulus sets^81,82^.

Neither differences in training regimes (supervised vs. self-supervised) nor in computational objectives (e.g., object vs action recognition) improved alignment with LOTC. While networks trained on action recognition did show some differences, such as a separated hand cluster compared to object-trained models (Figure 5d), they still failed to capture the action-related organization observed in LOTC. Why do models trained on action recognition do not show any better alignment with LOTC relative to standard object recognition models? One possibility is that the action categories used during training are too abstract. For instance, the label "opening" could refer to actions as different as opening a box or opening one’s eyes^43^, thereby failing to isolate action-effector relationships that drive LOTC responses. More generally, although these models are trained on short video clips, rather than static images, they process actions as static patterns across frames, lacking sensitivity to temporal dynamics, predictive processing, and temporal integration that humans naturally rely on^83^. Finally, human action perception is shaped not only by motion but also by social context and affordances^84^, factors that are entirely absent from current DANN models^83^. For instance, the comparison between DANNs and visual cortex is especially revealing when considering the case of shape: while both systems are sensitive to aspects of shape, such as elongation and aspect-ratio, shape information might be used for different purposes: exclusively for categorization in DANNs, where shape is indicative of category membership, and for more varied behaviorally-relevant goals in the brain, such as grasping, manipulation, and functional use of objects. This divergence may arise because DANNs are trained on passive visual tasks (e.g., classification), whereas biological vision is inherently linked to action planning and sensorimotor experience. A promising direction may involve training models through reinforcement learning in embodied agents, where tasks are grounded in action. For example, agents could learn to evaluate an object’s graspability or identify the specific parts relevant for grasping and functional use^85^ or learning actions in social contexts while interacting with humans^84^. Overall, while TDANNs represent a step forward in modelling visual cortex organization, we point to the necessity of using more ecological, varied tasks – beyond object or action classification – and the inclusion of biological constraints^86^ to fully model OTC object space (but see^87^).

In summary, this study demonstrates the critical role of the action dimension as an organizing principle of object representations in lateral occipitotemporal cortex. While artificial neural networks successfully replicated animacy-based organization, they failed to capture the action-based topography observed in the brain, despite their prominence in human functional organization. These findings underscore the importance of behaviorally relevant object properties in shaping the visual cortex’s topography and advance our understanding of how multidimensional representations support object vision in the human brain.

## Methods

### fMRI experiment and analyses

#### Participants

19 participants took part in the fMRI experiment (11 females, sex self-reported, mean age 25.6 years, standard deviation 6.06). One male participant was excluded due to head motion exceeding one voxel. All participants were right-handed except one, all had normal or corrected-to-normal vision, and no history of neurological disorder. All participants gave informed consent, and the Ethics Committee of the University of Trento approved the procedure.

#### Stimuli

The stimulus set included 6 categories (Figure 1). Part of the images were used in^35^. The set comprised 3 body-parts (hands, headless bodies, and faces), 3 inanimate object categories (tools, manipulable objects, and non-manipulable objects), and chairs as a control category. Each object category was associated with a different degree of action-related properties. Tools were defined as hand-held objects that are typically used to physically and directly act on another object or surface (e.g., hammer); therefore, tools are not only graspable and manipulable, but also serve as action-effectors, akin to our hands^33^. Manipulable objects are objects that can be grasped, lifted, and manipulated but are not usually used as action-effectors (e.g., glass). Finally, non-manipulable objects were defined as large objects that cannot be grasped nor manipulated (e.g., bed). To control for low- and mid-level visual features, the object categories were matched for their perceived shape and orientation (Figure 1). In addition, tools and manipulable objects were matched for real-world size, ensuring that any difference between the two categories cannot be attributed to their actual size. Three additional categories (monkey faces, headless monkey bodies, monkey hands) were part of the experimental design but are not analysed for this report. Each category included 12 grey-scale images with a white background of 400x400 pixels. Behavioral ratings confirmed that hands and tools were perceived as carrying the most action-related information, with mean scores of 6.3 and 5.7, respectively, on a 1–7 Likert scale. Specifically, hands were rated as conveying a higher level of action-related information than both bodies (4.5) and faces (3.4). Similarly, tools received higher ratings than both manipulable (3.3) and non-manipulable objects (2.9).

#### Scanning procedure

In the fMRI experiment we collected 8 runs per participant. Each run lasted 400 sec (200 volumes). Each image was presented for 0.4 s, with an ISI of 0.266 s, in blocks of 8 s (i.e., 12 images per block). For each subject and for each run, a fully randomized sequence of all conditions was repeated 4 times, with a fixation block of 16 seconds at the beginning, in the middle (between sequences), and at the end of each run.

Stimuli were presented with the Psychophysics Toolbox package^88^ in MATLAB (2021b) (The MathWorks). Images were projected onto a screen (8 x 8 degrees of visual angle) and shown to the participants through a mirror mounted on the head coil. Participants were instructed to fixate their gaze on the fixation cross in the middle of the screen and press a button whenever the same image was repeated twice in a row within each block. The repeating image appeared once per block. Behavioral performance during the task was quantified by calculating response accuracy (mean = 93%, SD = 2.7%) and reaction times (mean = 0.6 s, SD = 0.02 s) for hits. Accuracy was defined as the proportion of correctly identified target stimuli, with responses considered correct if made within two trials following the targets, taking into account the fast presentation of the stimuli (0.4 s) and the reaction time of participants.

#### Imaging parameters

The fMRI data was collected using a 3T Siemens scanner with a 64-channel head coil in the Center for Mind/Brain Sciences at the University of Trento. MRI volumes were collected using echo planar (EPI) T2*-weighted sequence, with repetition time (TR) of 2 s, echo time (TE) of 28 ms, flip angle (FA) of 75°, and field of view of 220 mm. Each volume contained 50 axial slices, covering the whole brain, with matrix size 200 x 200 mm and 3x3x3 mm voxel size. Slices were acquired with a multiband (multi-slice) sequence, with slice acceleration factor = 3. Anatomical images were acquired using the T1-weighted acquisition and MP-RAGE sequence, with a resolution of 1x1x1 mm.

#### Preprocessing

The preprocessing was conducted using the Statistical Parametric Mapping software package (SPM12, Wellcome Trust Centre for Neuroimaging London) and MATLAB (R2021b, The MathWorks). The following standard preprocessing steps were applied to functional images: spatial realignment (to the first image) to correct for head motion; slice-timing correction; coregistration of functional and anatomical images; normalization to a Montreal Neurological Institute’s ICMB152 template; and spatial smoothing by convolution with a Gaussian kernel of 4 mm FWHM^89^. Following exclusion criteria defined prior to preprocessing, runs in which the head movement exceeded the size of one voxel (in either translation or rotation) were excluded from subsequent analysis. Based on this criterion, we excluded one participant; additionally, we excluded five runs in total in three participants (two runs in two participants and one in another participant).

The preprocessed signal was then modelled for each voxel, in each participant, and for each condition using a general linear model (GLM). The GLM included 7 regressors of interest, one for each experimental condition, and 6 nuisance regressors corresponding to the 6 motion correction parameters (x, y, z for translation and rotation). Convolution of the haemodynamic response function with the boxcar function was used to model the predictors’ time course.

#### Vector-of-ROIs

To gain insights into the topographic organization of body parts and objects with different degree of action properties in left ventral and lateral occipitotemporal cortex (OTC), we used a vector-of-ROIs approach^18,90^. This analysis allows exploring, in an unbiased way, how the topographic organization of objects, characterized by different properties, changes along a large swath of cortex from lateral to ventral OTC. We focused on the left hemisphere, as tool selectivity is strongly left-lateralized and the hand-tool overlap is larger and more robust in the left hemisphere^32,91^ (see Supplementary Material for results in the right hemisphere). The vector-of-ROIs approach consists of the following steps: first, we defined two reference points (coordinates from^18^), located in a medial region in left ventral OTC (around the parahippocampal cortex [PHC]) and in a superior and posterior region in left lateral OTC (around the transverse occipital sulcus [TOS]). Then, we build a vector connecting the two points by fitting a spline. To make sure that the vector passes through anatomical landmarks relevant for their selectivity profile, we defined 6 anchor points based on coordinates from previous studies. Three were in the left ventral OTC: the medial fusiform gyrus previously shown to respond to tools (mFG^30^), the fusiform face area in the lateral fusiform gyrus (lFG^92^), and a region that responds to small objects around the occipitotemporal sulcus (OTS^17^); the other three were in the left lateral OTC: the anterior portion of the inferior temporal gyrus previously known to respond to small objects (aITG^17^), the hand-selective inferior temporal gyrus (pITG^32^), and the body-selective extrastriate body area within the lateral occipital sulcus (LOS^92^). After fitting the spline, along the vector, we generated a series of partially overlapping spheres of 6 mm with a distance radius of 3 mm. The beta values extracted from each sphere were employed to perform univariate and multivariate analyses. Furthermore, to investigate how each category-selective peak represents all object categories, we selected the activation peak in the vector-of-ROIs for all categories separately for ventral and lateral OTC and analysed their functional profile. Results were tested with two-tailed t-tests and corrected for multiple comparisons.

#### Category overlap analysis

We measured the amount of voxel overlap between the activation clusters for each condition, separately for ventral and lateral OTC. To do that, we selected two masks using a combination of functional and anatomical criteria; specifically, we used the Neuromorphometrics atlas (Neuromorphometrics, Inc.) to define regions within ventral and lateral OTC; ventral OTC included the fusiform gyrus and the parahippocampal gyrus, whereas lateral OTC included the inferior and middle occipital gyri and the inferior and middle temporal gyri; within these anatomical regions we selected all the active voxels with a contrast of all conditions vs. baseline with a liberal threshold (p < .05 uncorrected); these masks, which contain only the voxels modulated by visual information, were used for the subsequent analysis. To compute the overlap analysis, we calculated the number of active voxels within each of the two masks for each condition vs. all remaining conditions (e.g., hands vs. all others) with a more conservative threshold (p < .001 uncorrected at the voxel level and p < .05 FDR-corrected at the cluster level). Applying a cluster correction ensures that only contiguous voxels with a meaningful minimum size are considered for the analysis. The resulting active voxels were employed to compute the overlap index which was calculated pairwise for all possible combination of categories by taking the number of voxels common to two clusters (for instance, the voxels that are active for both hands and tools) and dividing it by the number of voxels of the smaller of the two clusters. An index of 0 indicated no overlap between two categories, whereas an index of 1 indicates that the smaller cluster of a category falls completely within the bigger cluster of the other category. Following previously adopted approaches (e.g.^93^), we calculated the overlap at the group level. Overlap analysis at the group level may introduce smoothing that overestimate the amount of overlap between categories; however, previous comparisons of overlap analyses based on single subjects vs. group analyses revealed little differences in the results between the two^94^; moreover, the use of relatively conservative thresholds and the use of selective contrasts ensure the control of overestimation of overlap effects.

#### Representational similarity analysis

From each sphere along the vector, we extracted the patterns of activation for each condition and correlated pairwise the patterns with each other to obtain a 6x6 correlational matrix. Values in the resulting correlation matrices represent how the pattern of activity for each category/stimulus correlates with the remaining categories/stimuli, allowing us to investigate how the representational space for the conditions changes from ventral to lateral OTC along the vector of ROIs. Representational similarity analysis (RSA^40^) was used to correlate (via Pearson) the matrix generated from each sphere along the vector-of-ROIs with three models capturing different properties of the stimuli: action, animacy, and aspect-ratio.

The action and the animacy models were generated based on ratings provided by an independent group of participants (n = 22, 13 females, mean age 23.3 years, SD = 1.96) that judged a subset of 36 stimuli, chosen randomly among the entire subset, using the inverse MDS procedure^95^. Specifically, to test action-effector properties, we asked participants to arrange the objects according to the degree to which an object or a body-part *“is typically used to physically/directly act on another object or surface”* similar to the definition used in^33^. To test animacy, we asked participants to “*arrange the stimuli according to their animacy properties”*. To measure the overall shape of objects, a formula that captures aspect-ratio was used to test the influence of visual features in explaining patterns of activations for the inanimate objects, as most tools are elongated objects as they must be grasped to fulfil their function. The model was generated by calculating the aspect-ratio for all 72 stimuli using the following formula (as in^12^):

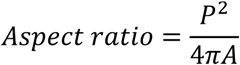

Where P is the perimeter of the object within the image and A is its area.

We generated the dissimilarity matrices for the models by computing pairwise the Euclidean distance between each value for each stimulus along the three dimensions. The three models are orthogonal to each other (see Results), indicating that they are independent and do not overlap in their predictions or dimensions. We calculated the lower bound of the noise ceiling by iteratively correlating each subject matrix with all the remaining subjects’ group-average matrix, leading to a final score that indicates the best possible fit to the neural data that the model can achieve given the noise in the data^96^. Confirming the high reliability of the data, the lower bound of the noise ceiling across lateral and ventral OTC ranged from 0.8-0.9 in VOTC and from 0.7-0.8 in LOTC (Figure 5b), indicating a strong correspondence across participants’ activity patterns.

#### Index analysis

The values of correlation matrices (as generated above) were used to calculate two indices: the grasp index and the action-effector index. These indices capture the degree to which the representational content of each body part’s activity pattern is correlated with the representational content capturing the action-effector and graspability properties of objects. The action-effector index measures the degree to which each body part relates to objects that are characterized as being action effectors, a property that is specific to tools (e.g., hammer) and not shared with other manipulable objects (e.g., we can grasp and manipulate a glass, but we do not typically use it to act on something else). The grasp index represents the degree to which each body part relates to objects that can be grasped and held in hands, a property common to both manipulable objects and tools (e.g., a glass and a hammer are both graspable), but not to large non-manipulable objects. To calculate the action-effector index, for each participant we took the correlation between each body-part with tools and from that we subtracted the correlation between each body-part and manipulable objects (e.g., body-tool minus body-manipulable). To calculate the grasp index, for each participant, we took the correlation between each body-part with manipulable objects and from that we subtracted the correlation between each body-part and non-manipulable objects (e.g., body-manipulable minus body-non-manipulable). All results were corrected for multiple comparison using Bonferroni correction.

### Deep Artificial Neural Networks

We tested a series of deep artificial neural networks (DANNs) to test the possible convergence or divergence in the topographic organization and representational profile between visual cortex and DANNs. We selected three different models varying in architecture and training task which are described in detail below.

#### Non-topographic networks

We selected two non-topographic networks based on the ResNet-50 architecture^97^ trained either in object recognition or action recognition. ResNet-object, trained in object recognition with ImageNet^42^, has been shown to effectively capture representations within category-selective areas in visual cortex^98^. ResNet-action, trained in action recognition with Moments-in-Time^43^, was chosen to test the influence of a training task focused on action recognition in capturing neural responses for action-related categories.

#### Topographic networks

As these standard networks do not have topographic constraints, we selected a further recently developed family of models that implement some constraints within their architecture to mimic the topographic organization of visual cortex^26^. These models – called Topographic Deep Artificial Neural Networks or TDANNs – were based on a ResNet-18 architecture and were trained with a self-supervised contrastive learning task^40^ on the ImageNet dataset. Prior to training, a mapping of units is implemented within each layer of the network, so that each unit has a corresponding 2D coordinate that maps them into a 2D grid that represents their physical distance. During training, a spatial loss function (together with the self-supervised task loss) is introduced: this function constraints nearby units to have correlated firing patterns to the same features within the dataset, so that the units that have similar functional properties will fall close in the simulated physical space. A parameter called *α* in the spatial loss function indicates how much the neighbouring units must be correlated with each other; following^26^, we used a value of *α* = 0.25, as it has been demonstrated to be the optimal value for the emergence of VTC-like topographic organization. These networks include 8 layers implementing topographic constraints, with different surface areas across layers to simulate the hierarchy of the ventral visual stream, from V1 to high-level VTC. We use five different random initializations of the network weights.

#### Data analyses

##### Univariate

For the TDANN only, we performed simulated “univariate” analysis by testing the topographic organization and selectivity profile of the five different random initializations of the network in response to our six object categories; most analyses were conducted on the last layer that qualitatively showed the clearest clustering by categories, which we called VTC-like layer (as in^26^). Specifically, we tested 1) the clustering of units selective for the different object categories within the simulated physical cortical space in the VTC-like layer and 2) the selectivity profile of the top-50 most selective neurons for each category in the VTC-like layer. *Overlap:* To examine whether object categories in the VTC-like layer of the TDANN exhibit a similar relationship to those found in the OTC, we measured the overlap in selectivity between units across different conditions. We followed the method introduced by^26^. Specifically, the simulated cortical sheet was partitioned into 1 mm wide square sections. In each section, we assessed the proportion of units that were selective (t > 3.5) for two categories (e.g., hands and tools, hands and faces, etc.) in pairs. The overlap between these categories was determined by analyzing the frequency of selectivity co-occurrence of the two categories within each section. Essentially, if the selectivity frequency for one category can predict the selectivity for the other, the unit populations are considered to overlap. This overlap is measured using an index that ranges from 0 to 1: a score of 0 means the presence of units selective for one category (e.g., hands) always predicts the absence of units selective for the other (e.g., tools); a score of 0.5 indicates no predictability between the two categories; and a score of 1 signifies perfect overlap, where the presence of units selective for one category always coincides with the presence of the other category.

##### Multivariate

For all networks, we presented our stimulus set and extracted the feature activations from the convolutional and fully-connected layers across the network hierarchy for the DANNs, and from the eight topographic layers for the TDANN. We generated RDMs for each layer by correlating pairwise the features extracted by the networks for each stimulus. As for neural data, for each layer in each network, we performed the RSA analysis testing three models (shape, animacy, and action) and computed the action-effector and grasp indices. Moreover, we computed multidimensional scaling on the matrix of the last convolutional layer of the two ResNet and of the VTC-like layer of the TDANN, to explore its multidimensional profile more in detail. Statistical significance for all results was assessed via 10000 permutation tests (p = .001).

## Data Availability

The following publicly available resources were used for this work: pretrained ResNet-action with Moments-in-Time and ResNet-object with ImageNet: https://github.com/zhoubolei/moments_models, TDANNs: https://github.com/neuroailab/TDANN.

All types of brain images and statistics data are available from the authors upon request. The data used for final statistics is available through the Open Science Framework: https://osf.io/ctmbx/.

## Code Availability

Matlab code used to analyze the data is available on the Open Science Framework at the following link: https://osf.io/ctmbx/

## Author Contributions

D.C., S.B., and H.O.B. designed the experiment.

D.C. and S.B. designed the experiment, interpreted the data.

D.C. collected and analyzed the fMRI data.

N.T. wrote and analyzed the code for deep neural networks.

D.C., S.B. and N.T. wrote the first draft.

All authors reviewed the paper.

## Supplementary Materials

### Supplementary Figures

**Figure S1.**
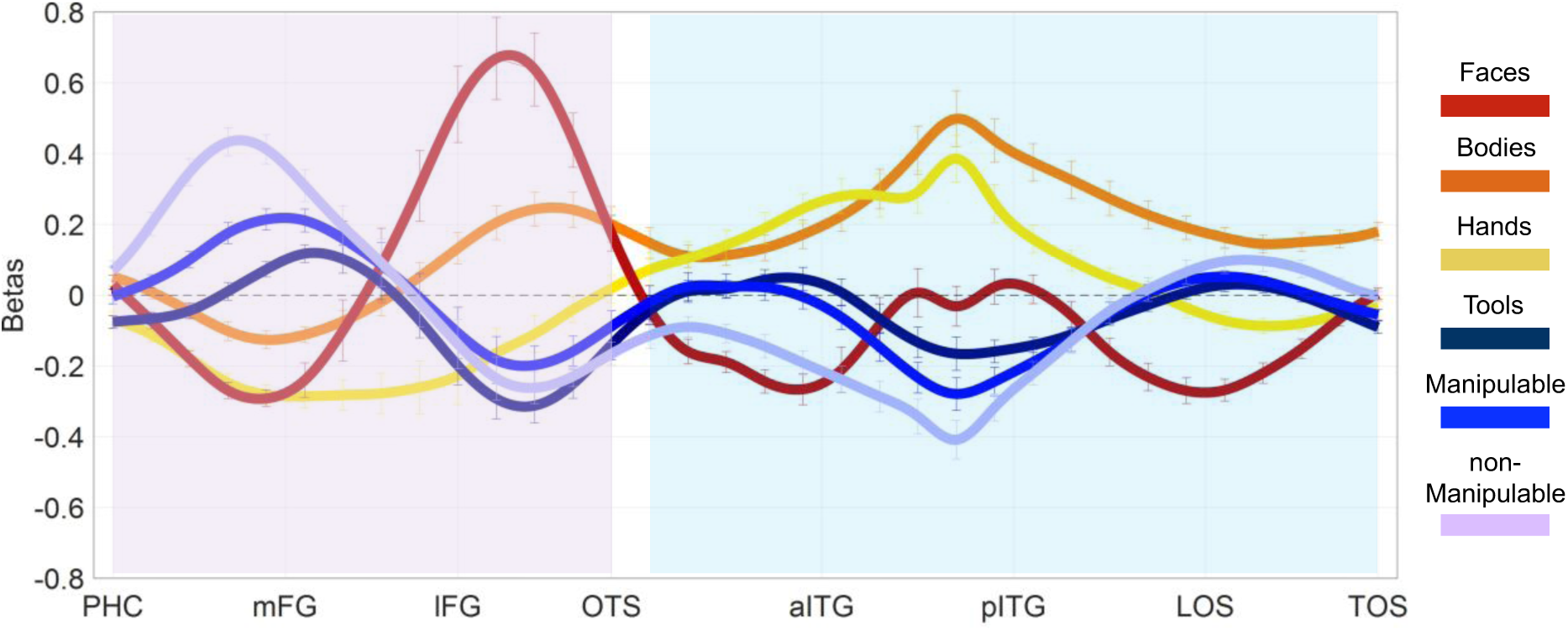
Right hemisphere vector-of-ROIs. The same procedure to generate the vector was followed as for the left hemisphere (see methods for details). The spheres along the vector cover an analogous portion of OTC as in the left hemisphere. Normalized activation (against the average of all categories) is plotted for each category as a function of the position of the vector along the cortex. The x-axis corresponds to each sphere along the vector, with labels for major anatomical landmarks; the Y axis corresponds to the normalized beta values. The vector was broadly divided into a ventral component (pink shade) and a lateral component (light blue shade). Contrary to the left hemisphere, no action-related organization can be observed in right lateral OTC. Error bars correspond to ± 1 SEM across subjects.

**Figure S2.**
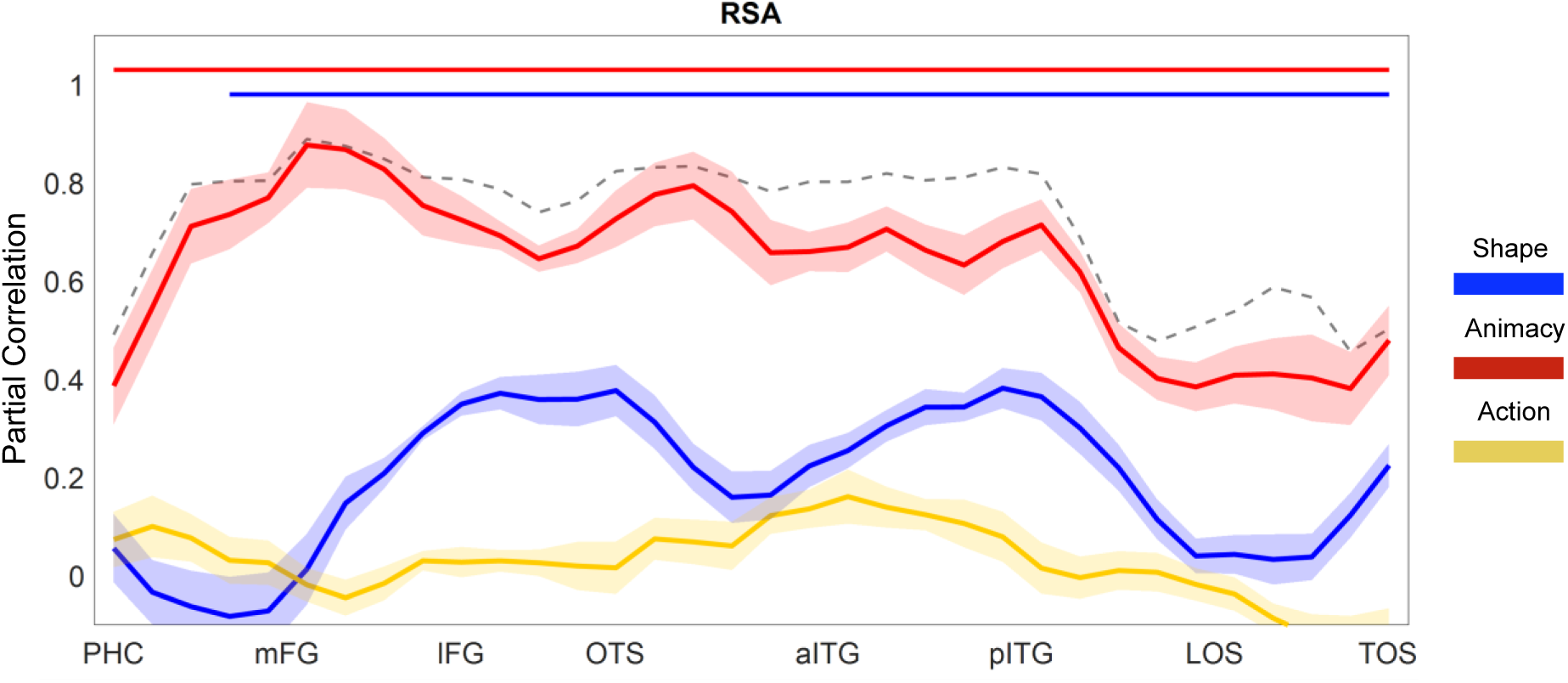
Object dimensions in the right hemisphere. Vector-of-ROIs RSA results. The dashed line represents lower bound of the noise ceiling. Horizontal lines indicate statistical significance (vs. baseline) for each model (*p* < .0013 Bonferroni corrected for n = 34 comparisons). In no sphere of the vector there is a significant effect for the action model, indicating that animacy and – secondarily – shape dominates the object space in the right hemisphere.

**Figure S3.**
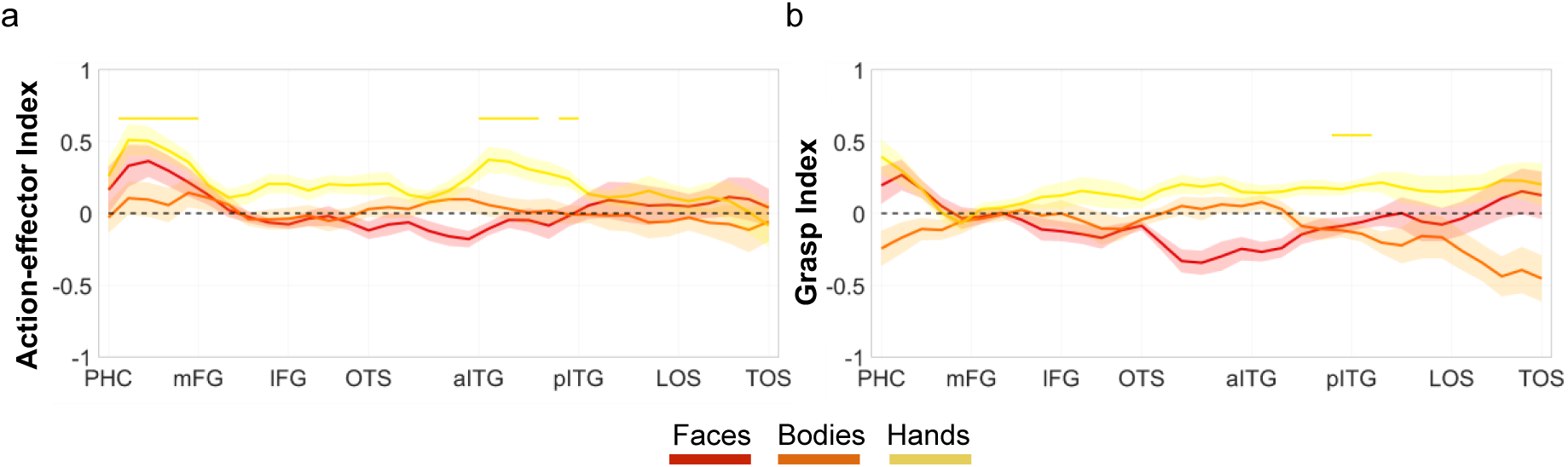
Index analysis. a) Vector-of-ROIs action-effector index. b) Vector-of-ROIs grasp index. Color-coded lines at the top of each plot indicate spheres along the vector where each index reached significance, corrected for the number of spheres (n = 34; p = .0015). Some effects can be found in right LOTC and VOTC, indicating that, despite the lack of the general action information, hand and tools are moderately correlated with each other also in the right hemisphere.

